# Simulation of receptor triggering by kinetic segregation shows role of oligomers and close-contacts

**DOI:** 10.1101/2021.09.29.462451

**Authors:** Rob Taylor, Jun Allard, Elizabeth L Read

## Abstract

The activation of T cells, key players of the immune system, involves local evacuation of phosphatase CD45 from a region of the T cell’s surface, segregating it from the T cell receptor. What drives this evacuation? In the presence of antigen, what ensures evacuation happens in the sub-second timescales necessary to initiate signaling? In the absence of antigen, what mechanisms ensure evacuation does not happen spontaneously, which could cause signaling errors? Phenomena known to influence spatial organization of CD45 or similar surface molecules include diffusive motion in the lipid bilayer, oligomerization reactions, and mechanical compression against a nearby surface, such as that of the cell presenting antigen. Computer simulations can investigate hypothesized spatiotemporal mechanisms of T cell signaling. The challenge to computational studies of evacuation is that the base process, spontaneous evacuation by simple diffusion, is in the extreme rare event limit, meaning direct stochastic simulation is unfeasible. Here we combine particle-based spatial stochastic simulation with the Weighted Ensemble method for rare events to compute the mean first-passage time for cell surface availability by surface reorganization of CD45. We confirm mathematical estimates that, at physiological concentrations, spontaneous evacuation is extremely rare, roughly 300 years. We find that dimerization decreases the time required for evacuation. A weak bi-molecular interaction (dissociation constant estimate 460 microMolar) is sufficient for an order of magnitude reduction of spontaneous evacuation times, and oligomerization to hexamers reduces times to below 1 second. This introduces a mechanism whereby CD45 oligomerization could be accessible to an engineered therapeutic. For large regions of close-contact, such as those induced by large microvilli, molecular size and compressibility imply a nonzero re-entry probability 60%, decreasing evacuation times. Simulations show that these reduced evacuation times are still unrealistically long, suggesting that a yet-to-be-described mechanism, besides compressional exclusion at a close contact, drives evacuation.

**Statement of Significance:** In the immune system, T cells sensing pathogens depends on a process called T cell receptor triggering. In this process, proteins on the cell surface undergo reorganization, including local depletion of large membrane proteins from the area surrounding the T cell receptor. Computer simulations of protein dynamics provide a means to investigate phenomena in greater detail than that afforded by experiments. However, even simulations present challenges, because tracking the motion and interactions of individual molecules is computationally expensive. Combining a rare event algorithm with spatial simulations, we show that biochemical and mechanical properties drastically affect depletion timescales, and thus receptor triggering. Quantitative understanding of these timescales will constrain hypothesized mechanistic models and could suggest new strategies for T cell engineering.

## Introduction

In the immune system, a key step involved in T cells sensing pathogens is a process called T cell receptor triggering. In this process, the receptor binds to an antigen being presented by another cell, shown schematically in Fig. 1A(left), and initiates an intracellular signaling cascade. The physical and biochemical mechanisms of T cell receptor triggering are not fully understood. A number of different models have been proposed, reviewed in [1, 2]. These models synthesize a variety of experimental observations and help further understanding of how T cells can achieve exquisite sensitivity to small amounts of antigen, while also discriminating antigen from self.

**Figure 1:**
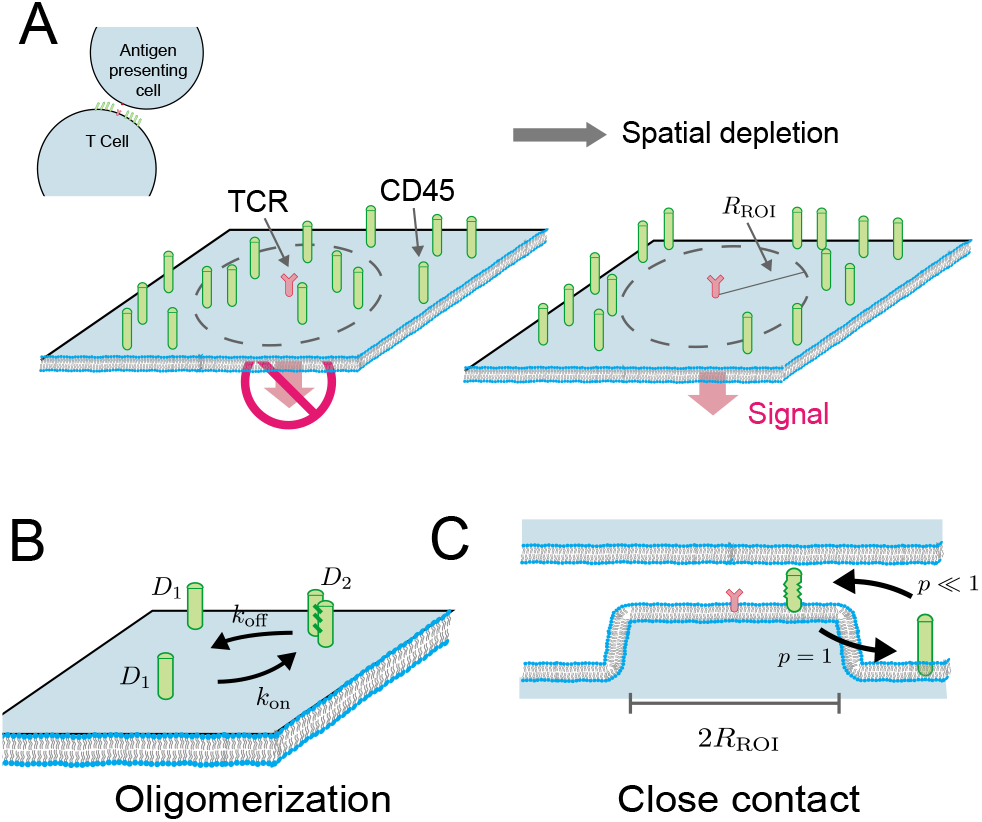
Surface dynamics for a large, membrane-bound surface molecule like CD45. (A) T Cell Receptor (TCR) binding to an antigen presented by another cell. (Left) If CD45 is uniformly distributed around the receptor, signaling is inhibited. (Right) The local depletion of CD45 from the region of interest (ROI) near the receptor, approximated here as a circle with radius *R*_ROI_, is a key step in T Cell receptor signaling [1, 2]. (B) Intermolecular interactions between CD45, e.g., oligomerization of CD45 into dimers. Diffusion coefficients for monomers and dimers are *D*_1_ and *D*_2_, respectively [13,14]. (C) Close contacts created by, e.g., microvilli, could lead to biased movement of CD45 due to compressional resistance of the molecule [15]. We represent these close contacts by probabilistically limiting entry into the ROI.

T cell receptor triggering is associated with reorganization of cell surface proteins, shown schematically in Fig. 1A. Local depletion of surface proteins with large ectodomains, such as the phosphatases CD45 and CD148, in the receptor’s vicinity has been demonstrated to be an important step [3, 4]. This depletion is consistent with a model of T Cell triggering known as the kinetic segregation model [1], in which large ectodomain proteins segregate from the close cell-cell contact necessary for receptor triggering, and depletion of these proteins from the receptor’s vicinity prevents them from interfering with the stable phosphorylation of cytoplasmic domains necessary to initiate the signaling cascade. The kinetic segregation model is supported by various lines of evidence: CD45 has been shown to have an inhibitory effect on receptor triggering [5, 6] and is found in low concentrations close to triggered receptors [6, 7], and synthetically holding a receptor in the CD45 depletion region augments signaling [8] The model is also consistent with the known geometry of the rigid extracellular domain of CD45, ~21 nm [7, 9], which is larger than the distance spanned by the receptor-antigen complex ~13nm [7, 10]. However, the kinetic segregation model cannot by itself fully explain T cell receptor triggering: other mechanisms likely contribute (e.g., see [1]) and CD45 plays somewhat contradictory roles [5].

How and when the local depletion (also hereon termed *evacuation)* of large ectodomain phosphatases from the receptor vicinity occurs remains unclear. It could happen pre-contact-formation (e.g., does local evacuation of CD45 clear the way for receptor-antigen binding?) or post-close-contact formation (e.g., a scenario where receptor-antigen binding is first enabled by close-contact due to a microvillus [11], or active membrane protrusion). The question of what drives this evacuation can be cast in three different lights: In the absence of receptor ligation, what mechanisms ensure evacuation does not happen accidentally? In the presence of ligation, what ensures evacuation happens in sub-second timescales necessary to initiate and sustain a signal? And, finally, if the process of evacuation tips the balance from inhibitory to stimulatory signaling in T cells, could modulating the evacuation process itself be an avenue accessible to engineered therapeutics?

Various mechanisms for this evacuation process have been proposed. These include: simple Brownian motion of CD45 in the plasma membrane [12]; oligomerization reactions between the molecules [13, 14] shown schematically in Fig. 1B; and mechanical compression by a nearby surface, such as that of the cell presenting the antigen. The compression region can be conceptually categorized as either a closecontact of ~100nm [15, 16], shown schematically in Fig. 1C, where there is no net lateral pressure on the molecules within the close contact, or something more similar to the wedge or tent shape resulting from a force on a single receptor pulling the membrane at a point [17]. The latter case has been studied theoretically [17, 18]. Beyond the scope explored in this work, there are many more possible mechanisms, including: spatial heterogeneity due to lipid composition or interaction with the cytoskeleton [19–21]; modulation of the configurational state of the individual molecules themselves can modulate their organization, e.g., by electrostatic interaction with the lipid membrane [22, 23]; or crowding out by CD3 [24].

Computer simulations of the spatiotemporal events involved in receptor triggering and immune synapse formation can provide a means to investigate phenomena that lie beyond the spatiotemporal resolution of measurement techniques [25]. In this paper, we investigate the evacuation process through computer simulations tracking reaction-diffusion dynamics of protein molecules on the cell surface. To simulate wide ranges of parameters, including different molecular phenomena, with a physiological and near-physiological numbers of molecules, we made recourse to the Weighted Ensemble algorithm [26], an enhanced sampling simulation method.

We confirm mathematical estimates that, at physiological concentrations, spontaneous evacuation is extremely rare [12]. We find that dimerization decreases the timescale of evacuation for even weak bi- molecular interaction by several orders of magnitude. The formation of higher-order oligomers reduces evacuation to a sub-second process, opening the possibility that an engineered oligomer of CD45 could significantly modulate receptor triggering. We find that formation of close contacts also decreases the timescale of evacuation. However, for large regions of close-contact, such as those induced by large microvilli, our model predicts evacuation times that are still too long by several orders of magnitude, using current estimates for the molecular size and compressibility of CD45. This suggests that the change in molecular motion driven by close contact alone is not sufficient to drive receptor triggering.

## Results

### A Rare-Event Reaction-Diffusion Simulation

Molecules in our model are represented as individual particles moving on a two-dimensional surface. Their mean density is *ρ*, and we assume there is a region of interest (ROI), for example near a single T cell receptor, that we approximate as a disk with radius *R*_ROI_. The quantity we wish to compute is the mean time until the ROI is empty, under various assumptions about the dynamics of the molecules – which we refer to as the evacuation time or mean first-passage time (MFPT). In the base case, we assume motion is purely diffusive with coefficient *D* (which could be thermal or include active, random forces [27]). Note that previous work [12] has shown that diffusion alone is too slow to be consistent with experimental data, giving MFPTs of ~ 10^10^ s whereas triggering can occur in reality within seconds [28, 29]. That is, the simultaneous evacuation of all molecules from the ROI by simple diffusion, given physiological surface density and ROI size, is a rare event. We are interested in the rare event limit because, first, its quantification helps reject this null hypothesis, and, second, it provides a necessary starting point for hypothesizing what may accelerate the biological process of T cell activation out of the rare event limit.

Assuming *D* ≈ 0.01 *μ*m^2^/s, and the radius of the ROI provides a characteristic lengthscale of 100 nm, then the characteristic timescale is 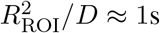. In these units, we estimate the surface density of CD45 to be *ρ* ≈ 9 molecules per 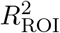[17].

#### Combining Weighted Ensemble and Smoldyn allows simulation of long-timescale stochastic spatial phenomena

To simulate evacuation in the rare-event limit, we combine the particle-based reaction-diffusion simulator Smoldyn [30] with a Weighted Ensemble algorithm [26, 31–33]. In this algorithm, shown schematically in Fig. 2A, many Smoldyn simulations are run in parallel, each assigned a weight *w_i_*, with ∑_*i*_ *W_i_* = 1. Periodic redistribution of weights and trajectories occurs after each time interval *τ*. More simulations are run in probabilistically less-likely regions of state space, but assigned a smaller weight. This focuses computational power on rare events while still maintaining an algorithmically exact statistical ensemble [32, 34]. Every *τ* time units, we measure the probability flux of trajectories into the evacuated state, shown in Fig. 2B. Trajectories that reach evacuation are killed, and their weights are redistributed into the system. Once the system reaches steady state, the mean flux (i.e., the summed weights of trajectories that reached the evacuated state) gives the reciprocal of the MFPT [31]. For full details see Methods and Supplemental Fig. S1. We name this algorithm and combination WE-Smoldyn.

**Figure 2.**
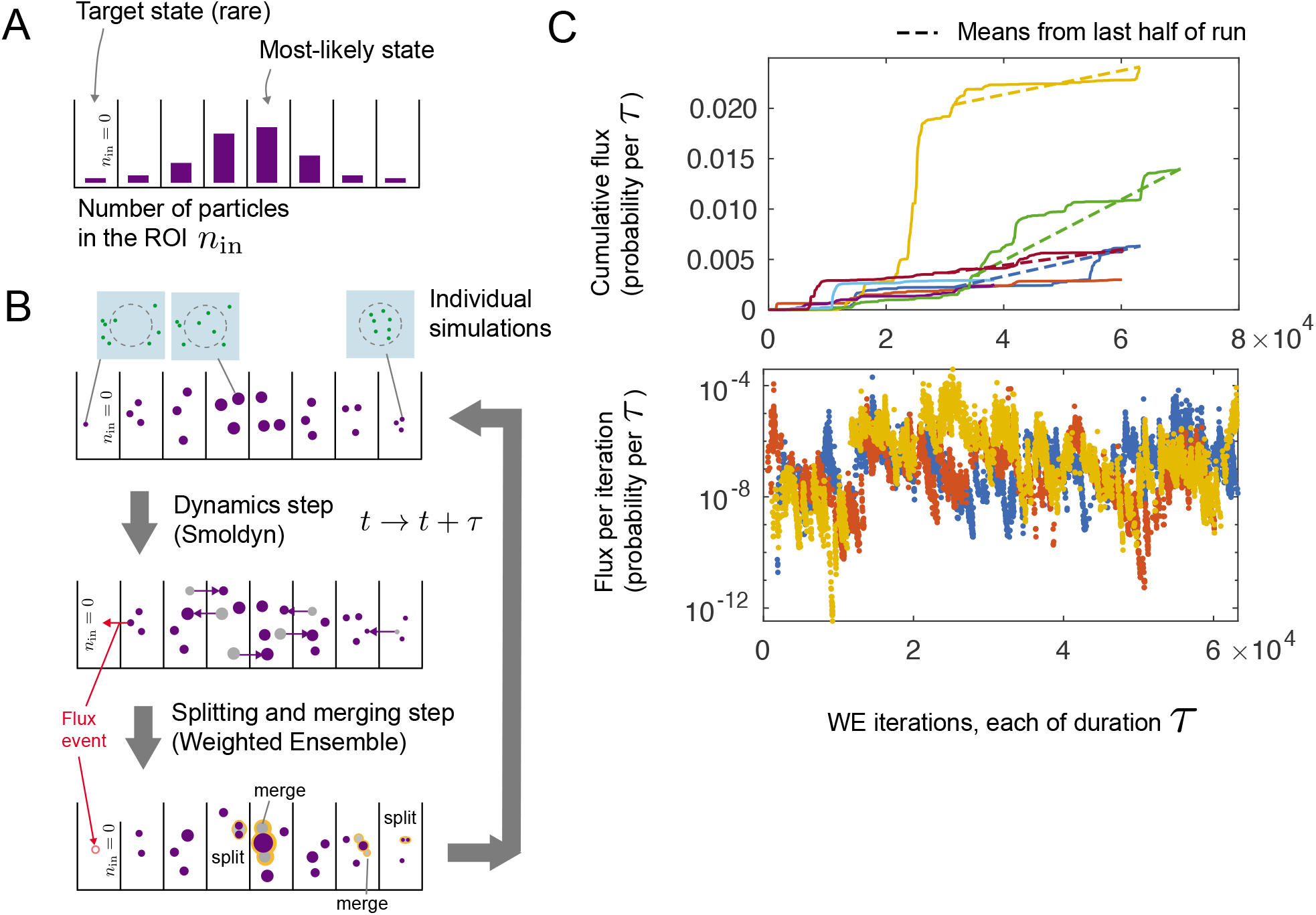
Weighted Ensemble algorithm with Smoldyn spatial dynamics simulations. (A) Partitioning of simulation state-space into bins based on the number of molecules in the ROI, 0 < *n*_in_ < *N*, where *N* is the total number of molecules. Bar heights represent the relative probability of the system to be found in the corresponding bin at equilibrium. In the simplest case in which molecules experience diffusion-only dynamics, this is a binomial distribution. (B) Description of the algorithm. Individual simulations (replicas) are shown as purple circles, and have statistical weights that can vary from replica to replica, represented here through circle size. Simulations are allowed to propagate according to dynamics simulated by the dynamics engine Smoldyn [30] for a period of time *τ*. After the dynamics step has completed, the replicas are reexamined to see their new bin locations. The number of replicas in each bin is then compared to *m*_targ_, the desired number of replicas in each bin. Bins with more than *m*_targ_ replicas have a “merging” event, where the replicas with the smallest individual weights are removed and their weight is redistributed to another replica within the same bin. Bins with fewer than *m*_targ_ (but still > 0) replicas have “splitting” events where the replicas with the most weight are duplicated into 2 daughter replicas, with the weight from the parent being redistributed equally to the daughters. Flux events, representing complete evacuations of the ROI, (red) have their replicas deleted and weight redistributed to replicas outside of the flux bin. For details on this redistribution of weights from flux events, see S1. (C) Flux measurements from unique runs displayed in two different ways: total flux accumulated (top, y-axis scaled linearly) and flux accumulated per WE iteration (bottom, y-axis scaled logarithmically; fewer runs shown for clarity). For each WE run, the measurements in the first half of the run are discarded to exclude the fluxes that might be measured during weight redistribution between the initial simulation state and the simulation state after many WE steps have passed. The mean flux measured during this period (slope of dashed lines) is then used to calculate mean first-passage time to the evacuated state. Cumulative fluxes for 7 runs are shown in the top figure, with the time series for 3 of those runs being shown in the bottom plot. Since the bottom plot is on a logarithmic scale, WE iterations where no flux was measured (in this case, ~ 90 % of iterations) do not appear.

#### WE-Smoldyn agrees with brute force stochastic simulation at low densities and approaches asymptotic calculation for high densities

We first simulate evacuation for a density of particles *ρ* undergoing diffusion only, and compute the mean time to evacuation. In Fig. 3A, we demonstrate the computational ability to simulate evacuation and compute MFPTs for a range of surface density values, reaching *ρ* = 10, which corresponds to a mean number inside the ROI of 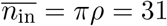. The MFPT grows superexponentially with *ρ*, reaching *T* ≈ 10^12^ seconds at the highest simulated density. The MFPT has an uncertainty of less than one order of magnitude (error bars are standard error of the mean).

**Figure 3:**
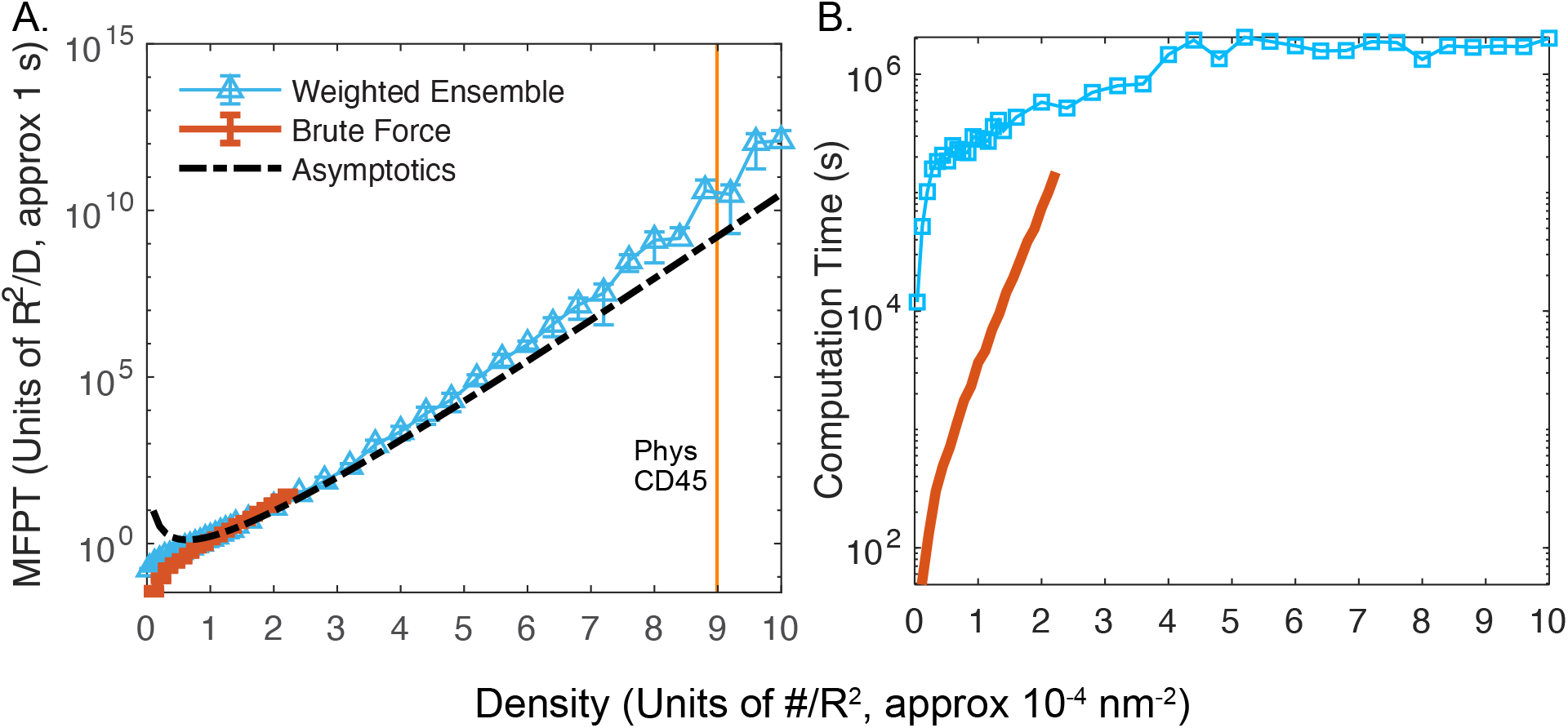
Mean evacuation time from the ROI for molecules only experiencing diffusion shows evacuation times well outside the experimentally observed timescales at physiological densities. Despite this drastic increase in evacuation times, use of Weighted Ensemble (WE) allows calculation on computationally-feasible timescales. (A) Evacuation times for densities ranging up to *ρ* = 10 molecules per 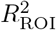. Estimate of physiological density of CD45 assuming an ROI of ~ 100nm in radius and ~ 9 × 10^-4^ molecules per nm^2^ (see main text) marked with a vertical line. Evacuation times from WE for domain size *L* = 5 (blue) agree with brute force simulations (red) at low *ρ* and with an asymptotic (infinite domain) approximation at low to intermediate *ρ* [12]. (Both the asymptotics and WE overestimate the evacuation times at very low *ρ*). Each WE data point uses 10 independent runs. Error bars for WE runs are computed by taking the average flux measured from the second half of each WE run as a single measurement, computing the 95% confidence interval of these flux measurements, and propagating these errors. Brute force error bars come from the standard error of the mean for 1000 runs. (B) Total computational costs for the data points in (A). WE data points come from summing the computation time of 10 independent WE simulations, with m_targ_ = 100 replicas per bin. WE runs were programmed to run until flux measurements from the final 1/3 of WE iterations and the middle 1/3 of WE met the following criteria: (i) Non-zero flux measurements in each third numbered at least 500 (ii) The KS-statistic between the two thirds reached a value of 0.02 or lower (iii) The KS-statistic between the two thirds with zeros removed reached a value of 0.3 or lower. Oftentimes, one or more of these requirements would not be met prior to reaching computation limits of an individual run, and in that case WE simulation would automatically stop after 2.5 days of computational time spent on WE and Smoldyn dynamics. Brute force data points come from summing the computation time of 1000 independent brute force evacuation events. Parameters were *τ* = 50Δ*t*, *m*_targ_ = 100, Δt = 10^-6^.

To validate our method, we compare with a brute-force simulation using Smoldyn, at low *ρ* (red). At high *ρ*, we compare to the asymptotic approximation from [12] (black dashed), which gives the following value for the MFPT

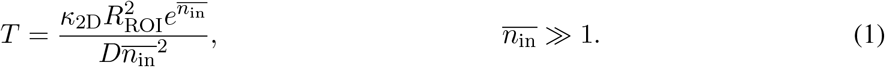

Here, *κ*_2D_ ≈ 0.7 is a constant independent of all parameters, see [12]. In the rare event limit, agreement to the asymptotics is within two orders of magnitude, and we hypothesize that the disagreement is due to a finite domain size in our simulation (whereas the asymptotic approximation is in the limit of infinite domain). We confirm this in Supplemental Fig. S2.

The computational scaling in Fig. 3B suggests that simple timestepping would take 4 × 10^5^ years of CPU time for 1000 evacuation events, whereas the Weighted Ensemble method took approximately 25 days per MFPT measurement (2.5 days per measurement, then repeated 10 times) on a single CPU core. Thus, the simulations we present throughout this paper would be unfeasible without recourse to an enhanced sampling algorithm like Weighted Ensemble.

The definition of evacuation time we use here and in [12] is instantaneous, in other words until the last molecule reaches the boundary of the ROI. This raises two notes: First, this is an approximation, since in the T Cell it is likely that a T cell receptor in an almost-evacuated ROI could still become triggered, just at a lower rate. Second, the timestepping algorithm we use here could lead to overestimates of the MFPT. To control for this second approximation, we confirm in Supplemental Fig. S2 that MFPTs are independent of simulation timestep Δ*t*.

### Oligomerization

#### Model for intermolecular interactions

There is some evidence that CD45 dimerizes [13, 14]. In this section, we explore the impact this would have on the evacuation process. To include dimerization, we add a reversible binding reaction to the model, with unbinding rate *k*_off_, and binding occurring whenever two particles are within a distance *r*_bind_ = 10^-2^, which corresponds to a physical distance of 1nm. Binding distance roughly corresponds to a binding rate, see Supplemental Fig. S3). We further assume that dimers diffuse more slowly by two-fold [35]. We explore the effect of varying *k*_off_, i.e., of varying the equilibrium constant for the dimerization reaction.

#### Dimerization decreases the timescale of evacuation by orders of magnitude even for weak bi-molecular interaction strengths

At low *k*_off_, evacuation times are decreased by over 5 orders of magnitude, as shown in Fig. 4A (towards left of horizontal axis). Order of magnitude changes in MFPT appear to track closely with corresponding steadystate monomer fraction, as a function of *k*_off_ (Fig. 4B). We can understand this heuristically as follows, making use of the asymptotic approximation (for monomer evacuation) from Newby and Allard [12]. At low *k*_off_, dimerization is dominant and most molecules are in dimer form. Evacuation is nominally slowed by the reduction in diffusion coefficient, since the evacuation time scales as *T* ∝ *D*^-1^ in the asymptotic (infinite domain) limit. However, this effect is outweighed by the reduction in the number of independent particles, since *T* ∝ e^*N*^/*N*^2^, which leads to an almost exponentially-lower MFPT [12]. Thus, the linear reduction in diffusion coefficient is dominated by the near-exponential dependence on the (linear) reduction in number of particles. Indeed when we compute MFPT as a function of the fraction of monomers in Fig. 4C, we observe an approximately exponential relationship.

**Figure 4:**
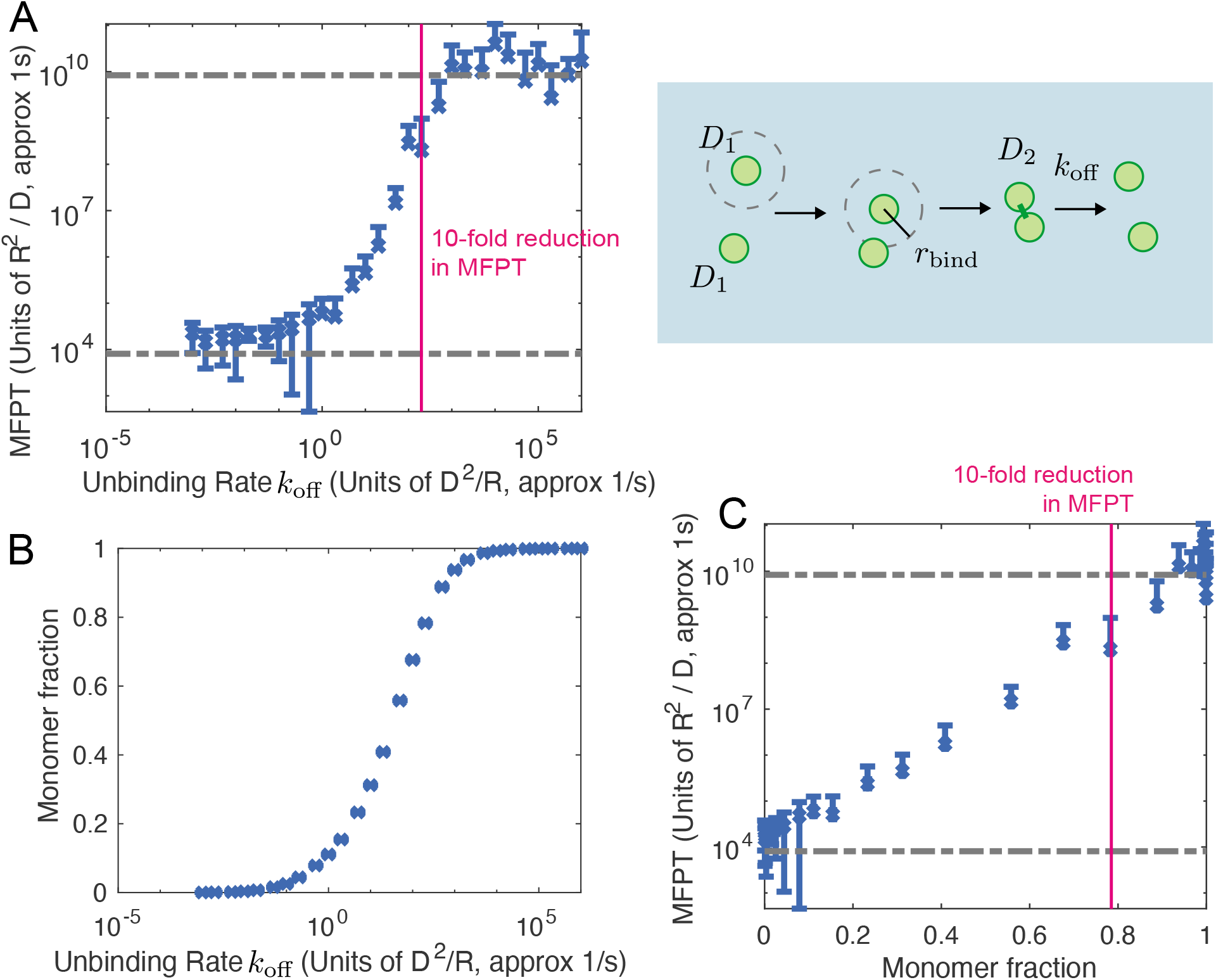
Dimerization of evacuating molecules causes several-order reduction in evacuation time (MFPT). (A) Right: Schematic of dimerization model used in Smoldyn. Molecules, diffusing at a rate *D*_1_, that get within a certain distance, *r*_bind_ = 0.001, of each other combine into a dimer that diffuses at a rate of *D*_2_ = *D*_1_/2. These dimers have an unbinding rate, *k*_off_, which varies across simulations. Left: Reduction in unbinding rate *k*_off_ (moving right-to-left on axes) leads to several-order reduction in evacuation time. At low dimerization fraction (high *k*_off_), evacuation time is the same as for non-interacting diffusing molecules with *ρ* = 9, *D* = *D*_1_, shown by the upper dashed line. At high dimerization fraction (low *k*_off_), evactuation time is the same as for non-interacting particles but with half the density and half the diffusion coefficient, i.e., *ρ* = 4.5, *D* = *D*_2_. (B) Average monomer fraction for the simulations given in (A) vs *k*_off_. (C) Same data as (A) and (B) showing the relationship between monomer fraction and evacuation time. A 10-fold reduction in evacuation time (vertical line in (A) and (C)) occurs when only 20% of the individual molecules are in a dimer, suggesting even weak binding can have drastic impacts on evacuation events.

More surprisingly, this dramatic reduction in evacuation time occurs even for weak dimerization. When monomer fraction is as high as 80%, meaning only 20% of CD45 subunits are in dimers, the MFPT is already reduced by ten-fold.

These relatively high unbinding rates correspond to weak homodimerization affinities. We can compute an effective 2D dissociation constant, defined as the concentration of reactants at which half the reactants are in the product (dimer). We find that a reaction with 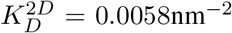 would yield a ten-fold reduction in evacuation time. A reaction with 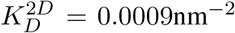 would yield a 1000-fold reduction. Conversion of 2D chemical properties to the equivalent 3D properties is nontrivial [8, 36], but a lower-bound estimate can be obtained by dividing the 2D density by the confinement height of the reaction [36–38]. In this case, the upper bound for the confinement height is the height of CD45 [9]. Using this, we can compute a lower-bound affinity for 10-fold reduction of MFPT, 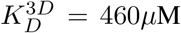, which corresponds to a standard binding free energy of 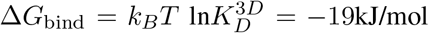. These binding strengths are an order of magnitude weaker than those measured for agonist TCR-peptide-MHC [39]. Note that these over-estimate the needed strength, since the confinement length we assumed to convert to a 3D affinity is an over-estimate.

#### Effects of higher-order oligomers on evacuation

This led us to wonder how evacuation times would be affected by the formation of higher-order complex molecular assemblies, for example as could be engineered using extracellular molecular linkers.

Full simulation of higher-order oligomerization was beyond the computational limits due to combinatorial complexity of number of molecular species. However, we can use Weighted Ensemble to compute the evacuation time assuming *k*_off_ is sufficiently low, such that all molecules are in the highest-order oligomer, shown in Fig. 5. Here, again, we assume that diffusion coefficient is reduced proportional to the number of subunits in the complex. For oligomers larger than dimers, we assume the ensemble is homogenous and only made of the largest complex. Between oligomer size 1 and 2, we show heterogeneous mixtures using the same sweep of *k*_off_ from Fig. 4, but plotted as a function of the average number of subunits in each independently diffusing particle,

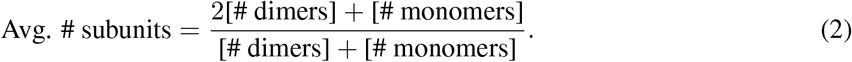

**Figure 5:**
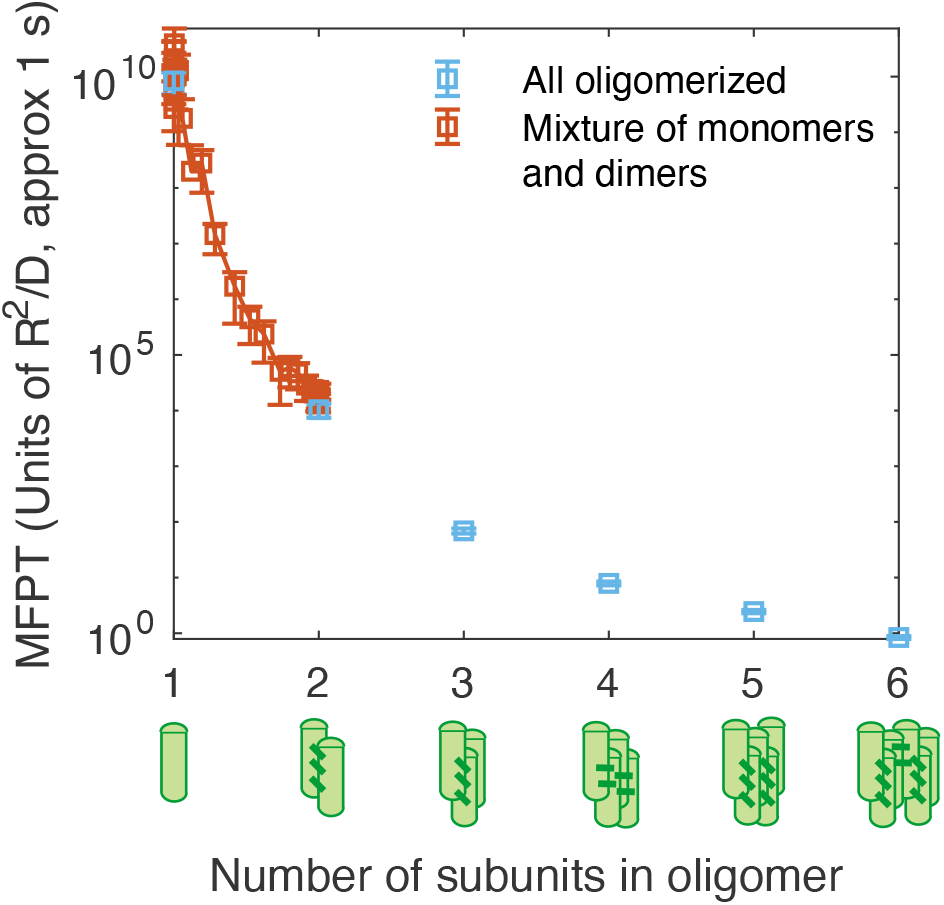
Formation of higher-order oligomers further reduces evacuation time, including times below ~ 1 second for hexamers. Assuming high-affinity (e.g., low *k*_off_) oligomerization would lead to homogenous ensemble of oligomers of a particular size (horizontal axis). Oligomer diffusion coefficients were taken to scale reciprocally with the number of subunits *D_n_* = *D*_1_/*n* [35]. Intermediate affinity oligomerization (red) would lead to heterogeneous monomeric and dimeric mixtures. We compute these intermediate evacuation times from dimerization data in Fig. 4 and Eq. 2.

Consistent with the result for dimers, these larger oligomers evacuate faster, despite diffusing more slowly. Indeed, hexamers with strong binding evacuate within 1 second.

### Close Contacts

#### Model for molecule behavior at a close contact

T cell receptor triggering can be induced by the formation of cellular protrusions called microvilli, which push against a surface, creating a region in close contact [15] between two surfaces, as shown schematically in Fig. 6A. If the close contact membrane-separation is smaller than the resting size of CD45, it has been hypothesized that this leads to dynamics in which CD45 can diffuse out but not back into the area of close contact [15]. Note that this is distinct from models in which the membrane deformation is tent-shaped or wedge-shaped, and therefore induces non-diffusive advection on compressed molecules [17]. We begin this section by exploring the model regime in which CD45 has an unspecified compressional resistance at a close contact size.

**Figure 6.**
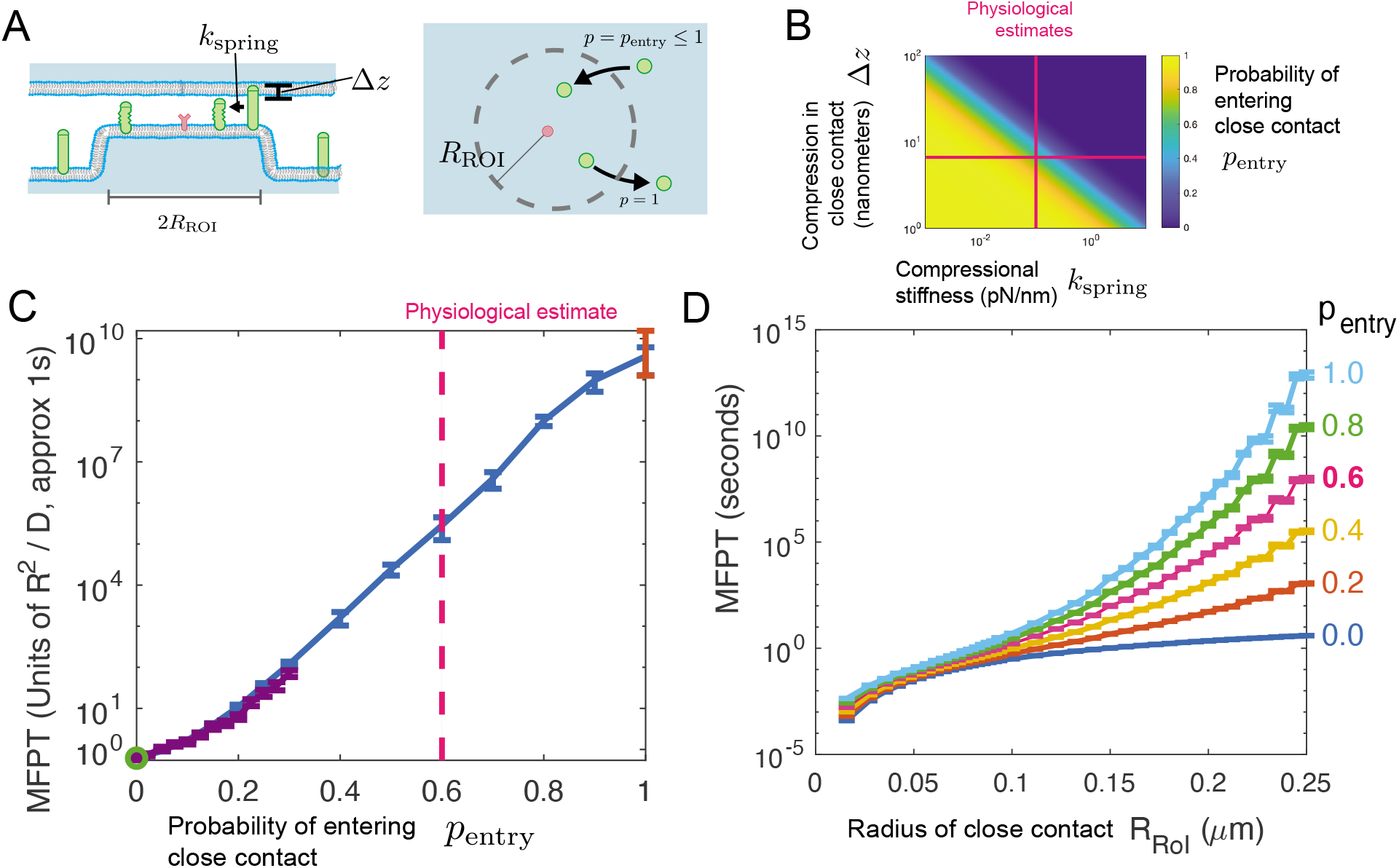
Formation of close contacts creates an energetic barrier for entry into the ROI. Even moderate energy barriers can create order of magnitude reduction in evacuation time at physiological densities. (A) Schematic of close contacts and how we chose to model them in Smoldyn. Close contacts, such as those from microvilli, can cause compression of large transmembrane particles such as CD45. The energy associated with this compression is modeled as a compressed spring *E*_spring_ = 1 /2*k*Δ*z*^2^, where *k* is the compressional spring constant, and Δ*z* is the distance of compression. This leads to dynamics in which movement into the ROI is reduced. We model the reduction as a probability penlry from the thermodynamic relation Eq. 3. (B) Heatmap of *p*_entry_ for a variety of compressional stiffnesses *k*_spring_ and the compression size Δ*z*. Pink lines provide reference for physiological estimates of both. (C) Evacuation time (MFPT) versus *p*_entry_, with the physiological estimate for *p*_entry_ given by the vertical dashed line. Included are results from WE simulations (blue) brute force simulations (purple). At *p*_entry_ = 1, we include the diffusion-only simulation result from Fig. 3 (red). At penlry = 0, we compute an analytic expression for the MFPT in Eq. 34 and Fig. S4, shown here in green. The inset shows, with a linear y-axis, close agreement between brute force and WE for *p*_entry_ < 0.1. For WE runs, an alternative method of redistributing weight in the flux bin is used, see S1. (D) Evacuation time versus radius of close contact, *R*_ROI_, in physical units (seconds, microns) assuming density of CD45 of 160*μ*m^-2^, assuming the ansatz Eq. 4.

Although the compressional resistance of CD45 is a key property in kinetic segregation models, estimates are challenging. Efforts to measure a similar molecule [40] have yielded estimates around *k*_spring_ = 0.1pN/nm. If a close contact is held in place by T cell receptors bound to antigen, the height difference between the rest size of CD45 and the close contact size has been estimated to be Δ*z* = 6.6nm [9].

#### For large regions of close-contact, such as those induced by large microvilli, molecular size and compressibility imply an intermediate re-entry probability

Using a Boltzmann relationship between the compression energy and the rentry probability (see Methods, Eq. 3), we can compute how these two molecular properties influence the ability of CD45 to enter the close contact, shown in Fig. 6B. Stiff or large molecules enter the ROI with near-zero probability, and soft or small molecules enter with high probability, but the physiologically estimated parameters lead to an entry probability of *p*_entry_ ≈ 0.6. Our finding is in contrast to previous models, e.g. [15], which assumed *P*_entry_ = 0. We perform simulations of the evacuation process with varying *p*_entry_ to investigate its effect on evacuation.

How well-approximated is the evacuation time by assuming zero re-entry, or by assuming free diffusion? It cannot be well-approximated by both, since the evacuation time we found above at *p*_entry_ = 1 is many orders of magnitude larger than the evacuation time in simulations from [15], who assumed *p*_entry_ = 0.

#### Physiological levels of molecular compressibility lead to significant reentry, leading in turn to significant delays in evacuation compared to purely one-way evacuation

We use our WE-Smoldyn algorithm to simulate a density *ρ* = 9 of molecules underdoing diffusion, but with the assumption that a given molecule, after exiting thte ROI, re-enters with probability *p*_entry_. We find that the evacuation times, shown in Fig. 6C, indeed vary between the simple diffusion case *p*_entry_ = 1 and the no-entry case *p*_entry_. The evacuation time, given physiological estimates of *k*_spring_ and Δ*z*, is around 10^5^ seconds (pink vertical bar in Fig. 6C). Although this value is orders of magnitude faster than the evacuation time computed for the *p*_entry_ = 1 case (simple diffusion), it remains substantially longer than T Cell triggering times.

We validate our results at *p*_entry_ = 1 by comparison with our simulations for simple, unhindered diffusion. We also solve for an analytic expression for the MFPT at *p*_entry_ = 0. This calculation is performed in the Supporting Text and shown in Supplemental Fig. S4. Agreement with WE-Smoldyn is shown as the open green circle in Fig. 6C. We further performed brute force simulation for 0 < *p*_entry_ < 0.3.

The dramatic effect of even small changes in *p*_entry_ led us to wonder about the relative importance of close contact size and gap size (which determined *p*_entry_). Note that, so far, all figure panels have shown evacuation time and densities nondimensionalized by scaling with the radius of the ROI. Rescaling to physical units, at a fixed physical density 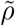, would require simulations over a range of *p*_entry_ and *ρ*. Instead, we make use of our finding that evacuation time is an approximately exponential function of *p*_entry_, and use this as an ansatz in Eq. 4 (Methods).

Evacuation times in physical units are shown in Fig. 6D, for a constant density 160 × *μ*m^-2^. Note this is lower than the physiological estimates by about 4-fold but at the limit of our current computational capability. At these parameters, a close contact region of radius less than ~ 100nm evacuates evacuates spontaneously in sub-second time. Close contacts that are perfectly impenetrable also evacuate in sub-second time, up to at least 250nm, comparable to the size observed [15, 41]. However, close contacts larger than 100nm with 20% or more re-entry probability have significant slow-down in evacuation. Here, an observed change in the shape of the curves leads to an interesting prediction: At low *p*_entry_ < 0.2, relative changes in *p*_entry_ lead to more signifcant changes in evacuation time compared to the same relative change in close contact radius. At high *p*_entry_ > 0.6, relative changes in close contact radius R_ROI_ lead to more significant changes in evacuation time compared to the same relative change in entry probability. At the physiological estimate *p*_entry_ = 0.6, roughly, a 20% reduction in *p*_entry_ has the same effect in reducing MFPT as a roughly 20% reduction in size of the close contact.

## Discussion

The paradigm of kinetic segregation — triggering a receptor by local depletion of its deactivating enzyme – has been proposed for a variety of signaling pathways [42, 43]. The most developed example is T cell receptor triggering by CD45 depletion. In this work, we show that, first, simple diffusive motion of CD45 leads to spontaneous depletion extremely rarely, in agreement with previous results [12]. Spontaneous depletion is therefore not at risk of false positive receptor triggering in the absence of an external cue. Second, we show that oligomerization of CD45 dramatically increases the speed of depletion. And third, we show that a close contact may accelerate depletion, but depending on its gap size and the mechanical properties of CD45, depletion may nevertheless be extremely slow.

Our results on oligomerization make a prediction: that externally-induced oligomerization of CD45 into higher-order structures would lead to more rapid receptor triggering, and indeed sufficient oligomerization (e.g., dominant heptamers, Fig. 5) would lead to spontaneous receptor triggering. Such oligomerization could be performed on the extracellular regions of CD45. Besides providing a test of the model predictions, this could provide a novel pathway for therapeutic modulation of T Cells in immunotherapies [44, 45]. The results thus provide a mechanism for how T Cell activity might be tuned by tuning CD45 oligomerization [13, 14].

Calculation of the theoretical evacuation time at a close contact has implications for models of close-contact surface molecule dynamics. In particular, Fernandes et al. [15] made the assumption that once a CD45 leaves the close contact region, it cannot re-enter (*p*_entry_ = 0). We confirm here that this leads to depletion times on the order of seconds. However, using estimates of CD45 geometry and mechanics, we compute that if there is a 60% chance of re-entry, the depletion time increases to 10^5^ seconds, much slower than observerd timescales of receptor triggering [15, 28, 29]. Formally, there are several possible resolutions to this discrepancy: If the estimated geometry and mechanical properties of CD45 are accurate, there must be another phenomenon driving evacuation. Alternatively, the molecular spacing could be smaller than estimated in [9], or the molecules could be much stiffer. Thus, the question of what mechanisms drive evacuation warrants further study.

The model we used is minimal in its assumptions and therefore subject to limitations. Our model focuses on the kinetic segregation mechanism, however, it has been proposed that kinetic segregation is just one of many mechanisms contributing to T cell receptor triggering [1]. Moreover, we focus on the inhibitory effect of CD45 on TCR signaling, whereas CD45 can both positively and negatively influence TCR signaling [5]. Within the kinetic segregation model, one limitation is our focus exclusively on total depletion, when the last molecule leaves the ROI. In reality, other steps in receptor triggering include ligand binding and receptor phosphorylation [16]. So, a more realistic model could be formulated in which the number of CD45 in the ROI determine a next-event rate. This rate would be high for total evacuation, and slower for partial evacuation. The MFPT one would study would be the time until the next-event has occurred. Without further assumptions, it is possible this next-even could happen slower than the total evacuation time, since it adds a subsequent step, or faster, since it can be triggered when evacuation is not total. Another limitation is the assumption of a flat, two-dimensional membrane. In particular, our consideration of microvilli ignored the purely geometric effect of a microvillus, in which distances around the perimeter of the microvillus are smaller than distances around the ROI in our flat simulations. Simulating diffusion on such curved surfaces is computationally challenging [46]. Yet another limitation is our focus on motion of CD45, when in reality the receptor moves as well [16]. Further integrative models, at the cell scale, may also include multiple receptors, and therefore multiple opportunities for a T cell to activate. Future research may explore these directions.

Crowding is prevalent in biology [47–53]. For that reason, there are examples in which un-crowding may be important — that is, when molecules must evacuate from a region before a given process can occur, and so the problem of making space is of general interest. These include the many transient cell-cell contacts which occur during tissue development (e.g., the delta-notch system [54,55]). There are also membrane-membrane contacts within cells, including between the endoplasmic reticulum and plasma membrane (where crowding could modulate interactions of molecules including Ora1 and Slim1 [56]). In 1D, an example is offered by transcriptional control in eukaryotes, which is achieved by the binding of many classes of proteins to DNA [57, 58]. Transcription factors (TFs) locate to binding sites within promoters and enhancers by 1D diffusion along the DNA and by attachment/detachment into the 3D cytoplasm [59–61]. The binding of larger structures, such as nucleosomes, which occupy ~ 150 base pairs (bp) of DNA, is inhibited by the presence of TFs, and therefore it is intriguing to wonder whether evacuation timescales are significant. Furthermore, enhancers, which are ~ 200 — 1000 bp stretches of DNA with 5-30 TF binding sites of various classes, may require evacuation of nucleosomes and transcriptional repressors to activate their target genes. Again in 1D, microtubules (inflexible polymers of the protein tubulin) are decorated by hundreds of microtubule-associated proteins [62, 63]. These proteins exhibit significant crowding [50, 51] and lateral diffusion along the microtubule lattice [64, 65]. Large microtubule-binding molecules may therefore have to wait for a region to be clear before binding.

Simulations performed here would be unfeasible without recourse to an enhanced sampling algorithm. Weighted Ensemble has been applied to many different types of stochastic dynamics simulations, however ongoing challenges are present, e.g., in *a priori* selection of state-space binning strategies, metaparameters, and weighting schemes to optimize convergence of desired observables [34]. Further systematic study of Weighted Ensemble metaparameter selection and analysis methods should lead to further increases in efficiency and empower future rare event simulation studies. Our WE-Smoldyn code base was built on top of the Smoldyn dynamics engine, which is widely used, flexible and with a large user base. We anticipate the combination with Weighted Ensemble and spatial stochastic simulation, as highlighted by full-featured software like MCell-WESTPA [66], will open new avenues of research, including for the evacuation questions posed in the previous paragraph.

## Methods

### Model and Dynamics

We represent the area surrounding the receptor as compartment with radius *R*_ROI_ in the center of a 2D square domain with edge length *L*, with the simplifying assumption being that the receptor motion is negligible relative to the motion of the individual CD45 molecules.

A summary of model variables, model parameters, Weighted Ensemble and simulation metaparameters is in Table S1.

#### Diffusion

Smoldyn is a time-stepping simulator with a continuous spatial domain (as opposed to a lattice-method). At each time step, molecule displacements are drawn from a Gaussian distribution whose width is determined by their diffusion coefficient *D*, with each chemical species having their own diffusion coefficient. At each Smoldyn timestep, Smoldyn tracks the location of each molecule and stopping when it observes a complete evacuation of the ROI (Fig. 1A, right). Aside from interactions with boundaries, barriers, and for molecular binding, all molecules diffuse independently and do not interact with each other.

As we are using a time-stepping based method, the determination of whether or not a molecule has evacuated the ROI within a timestep is based only on its starting and ending locations and specific details of the trajectory between the time-steps are lost. This representation results in evacuation events that occur between two sequential time-steps that are not observed by the method. As evacuation events are lost but none are gained, this would result in estimates of the MFPT that are higher than the true value rather than an underestimate. To minimize the number of evacuation events lost from these missing trajectories, we chose to use a small Smoldyn timestep, Δ*t* = 10^-6^. To ensure this choice of Δ*t* is small enough we confirmed that smaller timesteps give similar results, as shown in Fig. S2.

#### Intermolecular interactions and reversible dimerization

The diffusion coefficient for an oligomer is taken to be *D_n_* = *D*_1_/*n* where *n* is the number of CD45 subunits making up the oligomer, e.g. the dimeric diffusion coefficient is *D*_2_ = *D*_1_/2. Smoldyn uses an algorithm that is qualitatively similar to the Collins-Kimball model of bi-molecular reactions and approaches the Smoluchowski model for short time steps [67]. The association reaction occurs when two monomers diffuse within a pre-defined distance of each other referred to as the binding radius, *r_bind_*. Dissociation of dimers into two monomers is probabilistically determined at each time step, with probability determined by the detachment rate *k*_off_ [67].

Two-dimensional reactions are more complicated to analyze than their 3D counterparts [68, 69]. For example, there is no exact relationship between *r*_bind_ and a well-mixed *k*_on_. We confirm that, for our choice of Δ*t* = 10^-6^ and ranges of *r*_bind_ and *k*_off_, the steady-state unbound (monomeric) fraction is a smoothly increasing function of *k*_off_ and decreasing function of *r*_bind_, as shown in Fig. S3A. The dissociation constant, meaning the value of *k*_off_ at which half of subunits are in monomers, is a weakly increasing function of *r*_bind_, as expected by previous theoretical treatments [68, 69].

#### Close Contacts

By representing close contacts of the system as energy barriers caused from compression of molecules inside the ROI/close contact, we can then model these energy barriers by creating asymetric behavior between molecules attempting to enter the ROI and those leaving it. Molecules attempting to leave the ROI are free to do so, while those attempting to enter are only allowed to do so probabilistically. The energy barrier between the ROI and rest of the domain is taken to be the energy required to compress a spring *E*_spring_ = 1/2*k*Δ*z*^2^, where k is the physiological spring constant and Δ*z* is the size of the compression. The thermodynamic relationship between this probability and the energy compression is given by

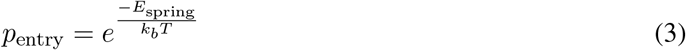

where *k_b_* is Boltzmann’s constant and *T* is the temperature. In the absence of evidence otherwise, we assume the presence of the close contact does not influence diffusion coefficient *D*.

#### Parameter estimates and model nondimensional scaling

The radius of the region of interest, *R*_ROI_, and the single-subunit diffusion coefficient *D* set a characteristic timescale of 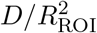.

Estimates for *R*_ROI_ range from 100—220 nm [15,17,41], depending in part on the definition, e.g., whether it is the minimum region necessary for receptor triggering, or the observed depletion zone size. The diffusion coefficient *D* has been estimated to range from 0.01*μ*m^2^/s [70] to 0.3*μ*m^2^/s [15]. Roughly setting *R*_ROI_ = 100nm and *D* = 0.01*μ*m^2^/s conveniently sets the characteristic timescale 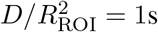.

Given the wide ranges of estimates, throughout this work we report times and distances in these scaled (nondimensional) units. Where appropriate, we report results in physical units, denoting these by explicitly including the unit (e.g. seconds or nanometers), and, if clarity necessitates, we use a tilde to denote the parameter with physical units. Domain size L and binding radius *r*_bind_ have units of *R*_ROI_. The corresponding physical parameters 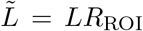 and 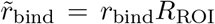 have units of nanometers or microns. The dynamics-engine timestep Δ*t* has units of 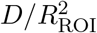 and the detachment rate *k*_off_ has units of 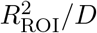.

The scaled molecule density *ρ* has units of molecules per 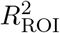, and the physical molecular density 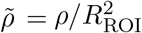 has units of molecules per square nanometer. Another interchangeable quantity is the number of particles in the ROI in a uniform distribution, 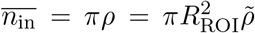. For CD45, estimates range from 482*μ*m^-2^ [15] to 1000*μ*m^-2^ [17]. For *R_roi_* = 100nm, this corresponds to a density ranging from *ρ* = 4.82 to *ρ* = 10. We use *ρ* = 9 as our focus [17] in all figures unless otherwise noted (e.g., in the *ρ* sweeps in Fig. 3 and the physical unit plot in Fig. 6D).

#### Ansatz for rescaling close contact evacuation time to physical units

To return to physical dimensions, a constant physical density 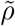 requires varying scaled density *ρ* since 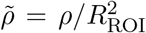. In Fig. 6D, we plot the evacuation time in physical units over a range of *R*_ROI_ Doing so would require a full exploration of both nondimensional *ρ* and *p*_entry_, which is outside of our computational capacity. So, as an approximation, we take the result in Fig. 6C, which suggests that evacuation time *T* (*p*_entry_) is a simple exponential function of *p*_entry_, that varies between the *p*_entry_ = 0 limit and *p*_entry_ = 1 limit. In other words,

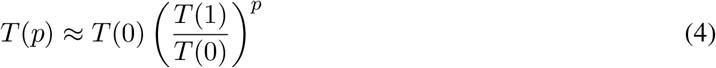

where *p* = *p*_entry_. The MFPTs for *T*(1) can be obtained from the simple diffusion simulations in Fig. 3A, while the MFPTs for *T*(0) can be obtained analytically, see Fig. S4B.

### Weighted Ensemble

Previous studies done on the evacuation of Brownian molecules in a 2D domain have found this evacuation to be a relatively rare event many orders of magnitude larger than typical time scales of the system [12]. A useful computational tool for studying rare events is the Weighted Ensemble sampling algorithm [26, 31,33] is statistically exact for many stochastic processes [32] and has been applied to many systems, especially in molecular dynamics [34].

#### Algorithm Overview

Weighted Ensemble obtains information of long timescale processes through multiple short timescale trajectories, hereafter denoted as replicas. A group of simulation replicas are initialized and attributed a probabilistic weight (see Fig. 2A). These replicas are allowed to evolve into a steady state distribution based on the ensemble space, which is then organized into bins based on location inside the state space. After the definition of these bins, the simulations are allowed to evolve for a fixed amount of time *τ* with periodic duplication and deletion of certain simulations (Fig. 2B. These duplications and deletions, dubbed splitting and merging, respectively, are done in such a way that preserves the total probabilistic weight of the system: splitting involves separating a simulation into two identical simulations each with half as much weight, and merging involves giving the weight from a deleted simulation to a simulation inside the same bin as the deleted simulation. While the total weight and its distribution between bins might change as the simulation evolves, the number of simulations inside each bin is manipulated so that computation power is evenly split between bins. Our binning order parameter is the number of subunits inside the ROI, n_in_. So each monomer inside the ROI increased the order parameter by 1, while for simulations with dimerization, each dimer inside the ROI increased the order parameter by 2.

If a replica reached the bin where *n*_in_ = 0, hereafter referred to as the flux bin, the replica is removed from memory, its probabilistic weight is recorded as outgoing flux, and then the weight is redistributed according to one of two different methods, see Fig. S1 and Reweighting Methods (below).

#### Model initialization

Each WE simulation is initialized with 1000 replicas of a Smoldyn simulation. In each replica, each of N molecules is randomly and uniformly placed throughout the entire domain. After this initialization, WE splitting and merging, and flux measurements are performed before each subsequent step of Smoldyn dynamics (Fig. S1C).

In simulations involving more than one molecular species, initialization is done with homogeneous molecular mixtures; either *N* monomers or *N*/2 dimers are uniformly distributed in the simulation, depending on which is closer to the steady state as found by brute force simulations in Fig. S3.

#### Reweighting Methods

There were two methods of redistributing weight removed from the system through flux into the flux bin (*n_in_* = 0, see Fig S1). The first method, which was the method used for all simulations not involving close-contacts, involves redistributing the weight by renormalization; the weight of all remaining replicas is scaled by the total weight remaining in the simulation. If replica *i* evacuates, it is removed and the weight for a replica *j* remaining in the simulation will scale according to

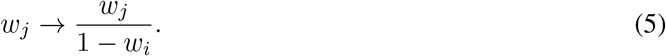

This method works for calculating the mean transition time from the steady-state distribution (or small perturbation from steady-state) to a rare fluctuation. In other words, we are measuring the MFPT from *A* → *B*, where *B* is defined as the bin *n_in_* = 0 (which is rarely visited), and *A* is defined as encompassing all bins *n_in_* > 0. Note that in many WE applications, significant time is required for the system to reach steady state, before which accurate MFPT estimates cannot be obtained from the averaged flux-to-target [31]. In our system, we know *a priori* the equilibrium distribution of particles undergoing simple diffusion. Our initialization of replicas according to the equilibrium distribution thus starts close to the non-equilibrium steady-state distribution (reached after some number of τ iterations), where the small weight entering the flux bin is continuously removed and returned to the remaining bins.

In close-contact simulations where *p*_entry_ ≈ 0 (Fig. 6), the steady state is the completely evacuated state. Here, we are seeking to compute a different transition time: from the uniform steady state (as if *p*_entry_ = 1) to the evacuated state, but where particles are restricted from re-entering the ROI. So, a second reweighting method was used for close contact simulations (see Fig S1B). In these simulations, weight from an evacuated replica is not redistributed to remaining replicas. Rather, each time a replica evacuates, a new replica is initialized as described above and given all of the weight from the evacuating replica. This ensures the the weight distribution throughout the state space remains statistically accurate, even when the flux of weight throughout the space is unidirectional.

#### Metaparameters

The above-described Weighted Ensemble method requires the specification of metaparameters *τ, m*_targ_, and the max number of iterations. In principle, the selection of these metaparameters should not impact the results of the WE simulation, but will impact the efficiency of convergence to an accurate MFPT.

In an effort to maximize the observed number of flux events, *m*_targ_ was chosen to be high to maximize the number of replicas in bins nearby the flux bin (transient bins), but not higher than allowed by computer memory limitations. There values were *m*_targ_ = 100 for Fig. 3, Fig. S2, and 200 for Fig. 4, Fig. 5.

The WE step was chosen to be *τ* = 50Δ*t*, which we found to be large enough to give replicas time to change bins before the splitting and merging process began, while also avoiding being too long to ensure a high number of replicas inside the transient bins.

### Implementation

Smoldyn simulations were executed through Smoldyn’s C library, libsmol. Combination of Smoldyn with Weighted Ensemble was written in C and is available to the public at https://github.com/dydtaylor/LibsmolWE. Execution of LibsmolWE simulations was done on UCI’s high performance cluster. Brute force Smoldyn simulations were executed in Libsmol, but outside of the LibsmolWE weighted ensemble framework. All data was analyzed in MATLAB. Evaluation of analytical solution for *p*_entry_ = 1 was done in Wolfram Mathematica.

### Analysis

To allot for burn-in time, i.e., an initial transient while the replicas approach a steady state, the first half of each run is discarded, and the mean flux measured in the second half of the run is taken to be a single measurement of the mean flux 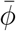, averaged across WE steps. Each WE data point in this paper is calculated from 10 independent WE simulations. These 10 repeats are then arithmetically averaged to give an estimate of the mean flux, averaged across repeats, 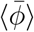, with error bars given by the standard error of the mean for the 10 WE simulations.

The average flux recorded from replicas reaching the flux bin, 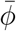 is used to calculate the MFPT from the Hill relation [31,71], 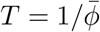. The error bars of the flux δφ then propagate to MFPT error bars by *δT* = |*T*|^2^*δφ*.

To calculate MFPT from brute force simulations, the arithmetic mean of 1000 (Fig. 3) or 500 (Fig. S4B) independent repeats was used, with error bars giving the standard error of the mean for those simulations. To calculate the single molecule first passage time distribution for Fig. S4A, an empirical CDF was created from the result of 20,000 brute force evacuations of a single molecule placed uniformly within the ROI.

To calculate monomer fractions from brute force simulations, purely monomeric Smoldyn simulations are initialized and run for 15 units of time. Afterwards, the monomer fraction we report is average measured over these last 5 time units.

### Method validation

Several methods were used to validate the WE results. For MFPTs where achieving brute force results was computationally viable, brute force results were included along with WE results (see Figs. 3, 6C). Monomer and dimer fractions were verified with brute force Smoldyn simulations (Fig. S3). For each binding radius presented, the sigmoidal monomer fraction vs unbinding rate curve was executed for a range of timesteps to verify the timestep of Δ*t* = 10^-6^ was small enough.

Close-contact WE simulations were found to agree with brute force simulations for low *p*_entry_ (Fig. 6C, inset). For *p*_entry_ = 1 we verified the endpoint with our simple diffusion WE simulations for *ρ* = 9.

An analytical solution for close-contact simulations when *p*_entry_ = 0 was obtained, see Supplemental. The analytic solution for a single molecule’s first passage time distribution was compared with an empirical CDF obtained from 20000 brute force Smoldyn simulations (Fig. S4A) and the analytical form for the MFPT for a variety of evacuating molecules was compared with the results from brute force Smoldyn simulations, 500 for each data point (Fig. S4B). The analytic solution for the parameters used in the close-contact WE simulations is included in Fig. 6C in both the main plot as well as the inset.

An asymptotic solution for the MFPT for homogeneous monomeric solutions was obtained from previous work [12]. When applicable, these asymptotics were used to verify WE results (Figs. 3A, S2). However, as can be seen in figures 3, S2, agreement with the asymptotics at higher densities is dependent on the size of the domain used. For a range of densities, we did a sweep of domain sizes (Fig. S2). Weighted Ensemble estimates reach to within one order of magnitude of the asymptotics above ≈ *L* = 5 at the highest simulated densities.

Convergence in time-step was done by comparing MFPT estimates for larger time-steps with the chosen time-step to ensure MFPT estimates did not undergo drastic differences in MFPT estimates. The time step of Δ*t* = 10^-6^, used for all simulations unless otherwise stated, is compared with with the timestep of Δ*t* = 5 × 10^-6^ is shown in supplemental figure S2B-D.

## Author Contributions

JA and ER designed research; RT developed code, performed research, and analyzed data; RT, JA, and ER wrote the manuscript.

## Acknowledgements

We thank Omer Dushek (Oxford), Jay Newby (U Alberta), Brian Chu (UC Irvine), and Dhiman Ray (UC Irvine) for valuable discussion. This work was supported by: NSF grant DMS 1715455 to ELR; NSF CAREER grant DMS 1454739 to JA; NSF grant DMS 1763272; and a grant from the Simons Foundation (594598, QN).

## Supplemental material

**Figure S1:**
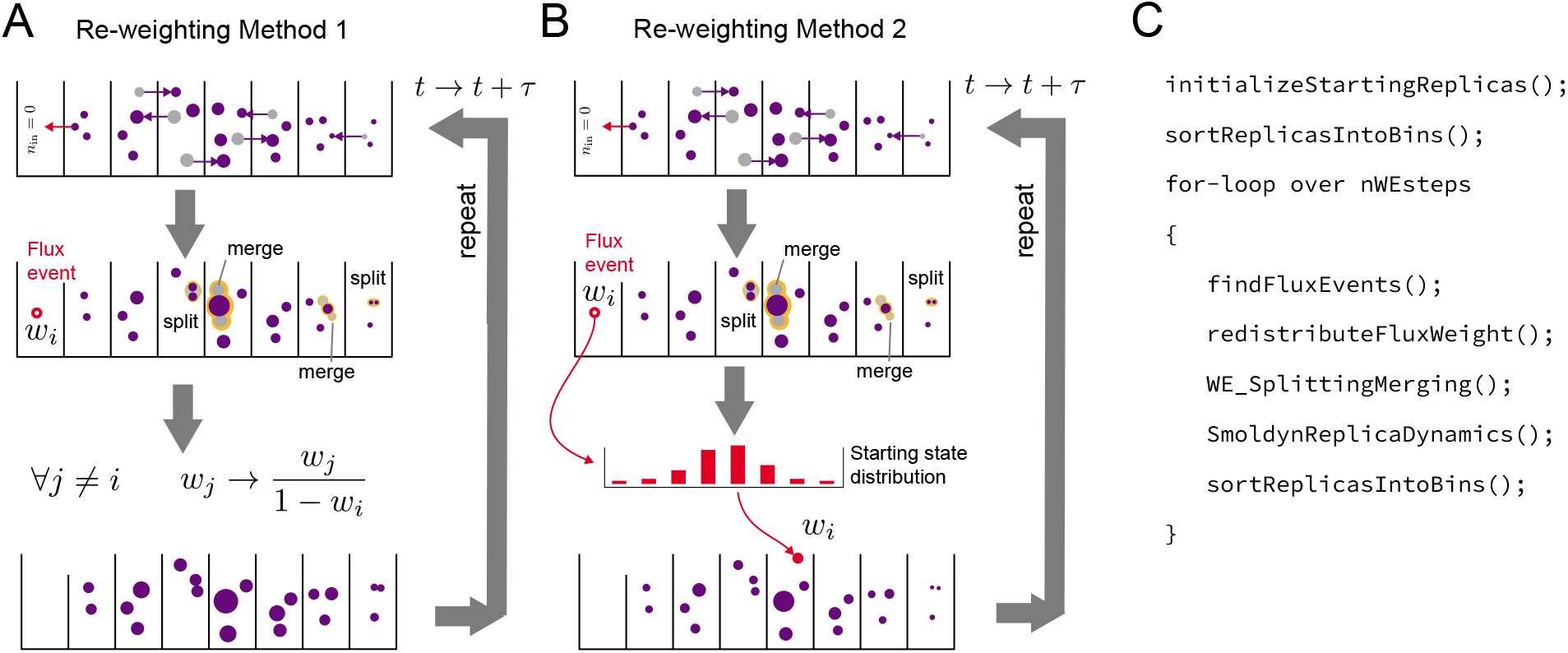
Different reweighting methods used for WE simulations. Method A is used for figures 2C, 3, 4, and 5. Method B is used for the WE runs in 6C and D. (A) Upon occurrence of a flux event, the weight from the flux event, *w_i_* is returned to the simulation by scaling the weight of the remaining simulations by their remaining weight, *w_j_* → *w_j_* /1 — *w_i_*. Note how weight distribution changes if replica movement between the bins is limited, e.g. in figure 6 with *p*_entry_ = 0. As bins are based on the number of molecules inside the ROI, no weight is allowed to reenter a bin after it leaves, which has a significant impact on steady-state weight distribution. (B) Upon occurrence of a flux event, the weight is given to a newly initialized replica drawn from the starting state distribution (all molecules are placed randomly and uniformly throughout the domain). (C) Pseudo-code of WE algorithm described in 2, including re-weighting method “redistributeFluxWeight()”.

**Figure S2:**
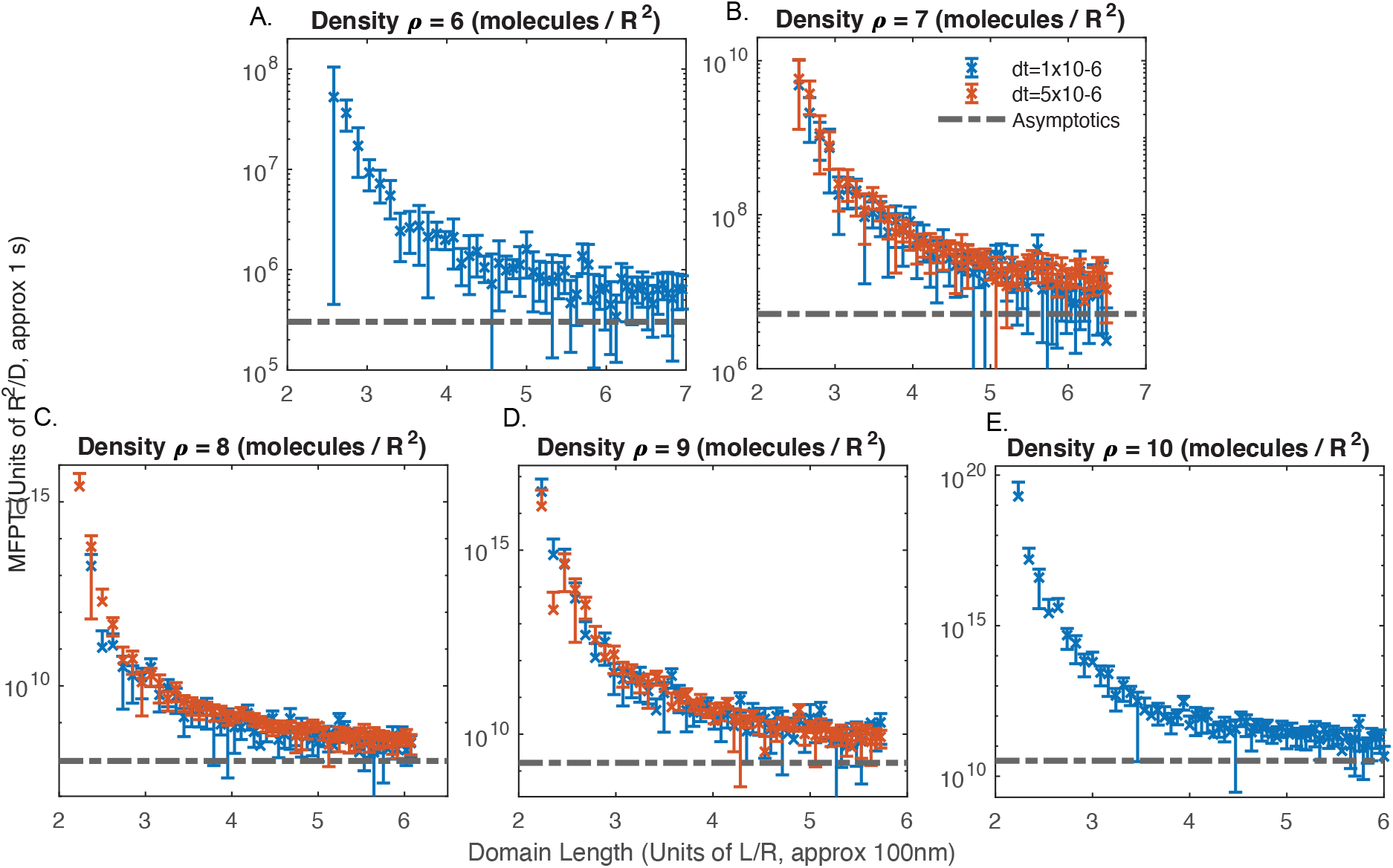
Evacuation time (MFPT) dependence on domain size *L* for given densities. (A-E) MFPT versus *L* for increasing values of density *ρ*. Horizontal dashed lines give the value of the infinite-domain limit from [12]. Weighted Ensemble estimates of evacuation time approach the asymptotic estimate as L increases. At density *ρ* = 9 (D), the Weighted Ensemble estimate reaches to within about one order of magnitude of the asymptotic near domain size *L* = 5. In light of this, dimerization simulations in the main text were chosen to have a domain size of *L* = 5.333 *R*_ROI_ (leading to total number of monomeric subunits *N* = 256 at *ρ* = 9). To investigate impact of time step on convergence, time steps of Δ*t* = 10^-6^ and Δ*t* = 5 × 10^-6^ were used for panels B,C and D. For all runs included, WE parameters were *τ* = 50Δ*t, m*_targ_ = 100. Each data point had 10 WE runs, with error bars representing standard error of the mean as calculated in Methods. Error bars were calculated from the 95% confidence interval of the mean flux, and propagating this error to the MFPT.

**Figure S3:**
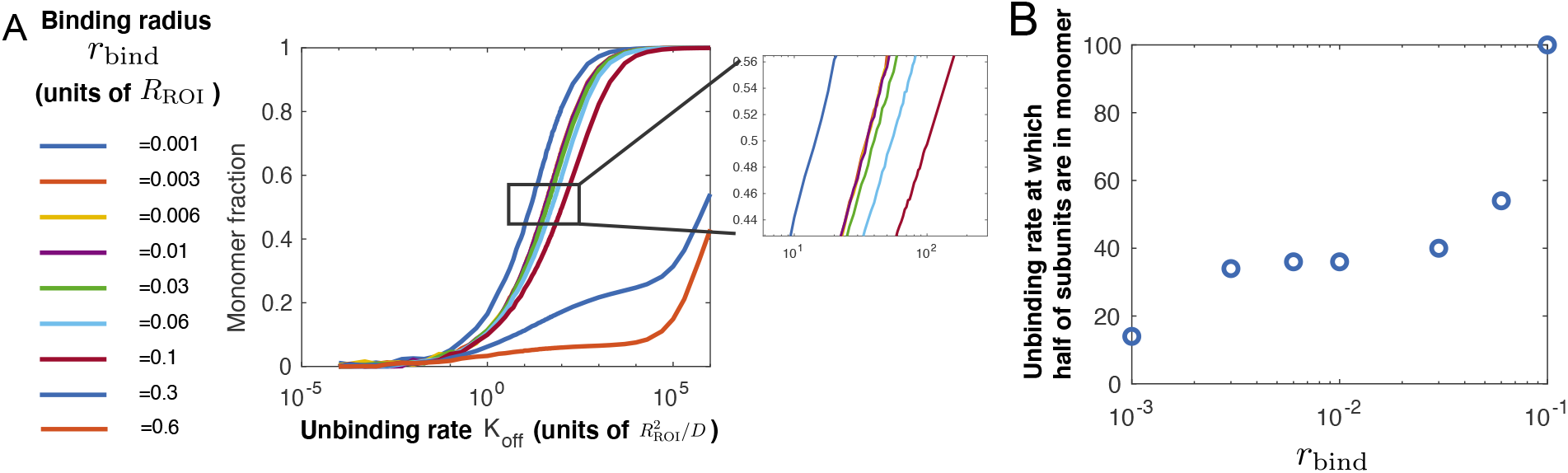
Particle-based reaction diffusion simulations of reversible binding kinetics on a 2D domain. Simulations were done in Smoldyn with 256 monomeric subunits in a square domain with side length L = 5.333 and run until reaching a steady state fraction of monomers. Each curve was independently found to be insensitive to decreasing the simulation time-step Δ*t*. (A) Monomer fraction at steady state for a range of *k*_off_ and *r*_bind_. (B) The effective dissociation constant, i.e., the unbinding rate *k*_off_ for which 50% of the monomeric subunits are in the monomeric form, as a function of binding radius. For binding radii close to the estimated physiological binding radius of *r*_bind_ = 10^-2^ (corresponding to roughly 1nm), the steady-state monomer we see weak insensitivity between the monomer fraction and the binding radius, as previously reported, due to the complicated logarithmic nature of two-dimensional diffusion [72, 73]. At binding radii larger than 0.3, the system approaches the crowding regime, resulting in large impact to monomer fraction.

**Table S1:**
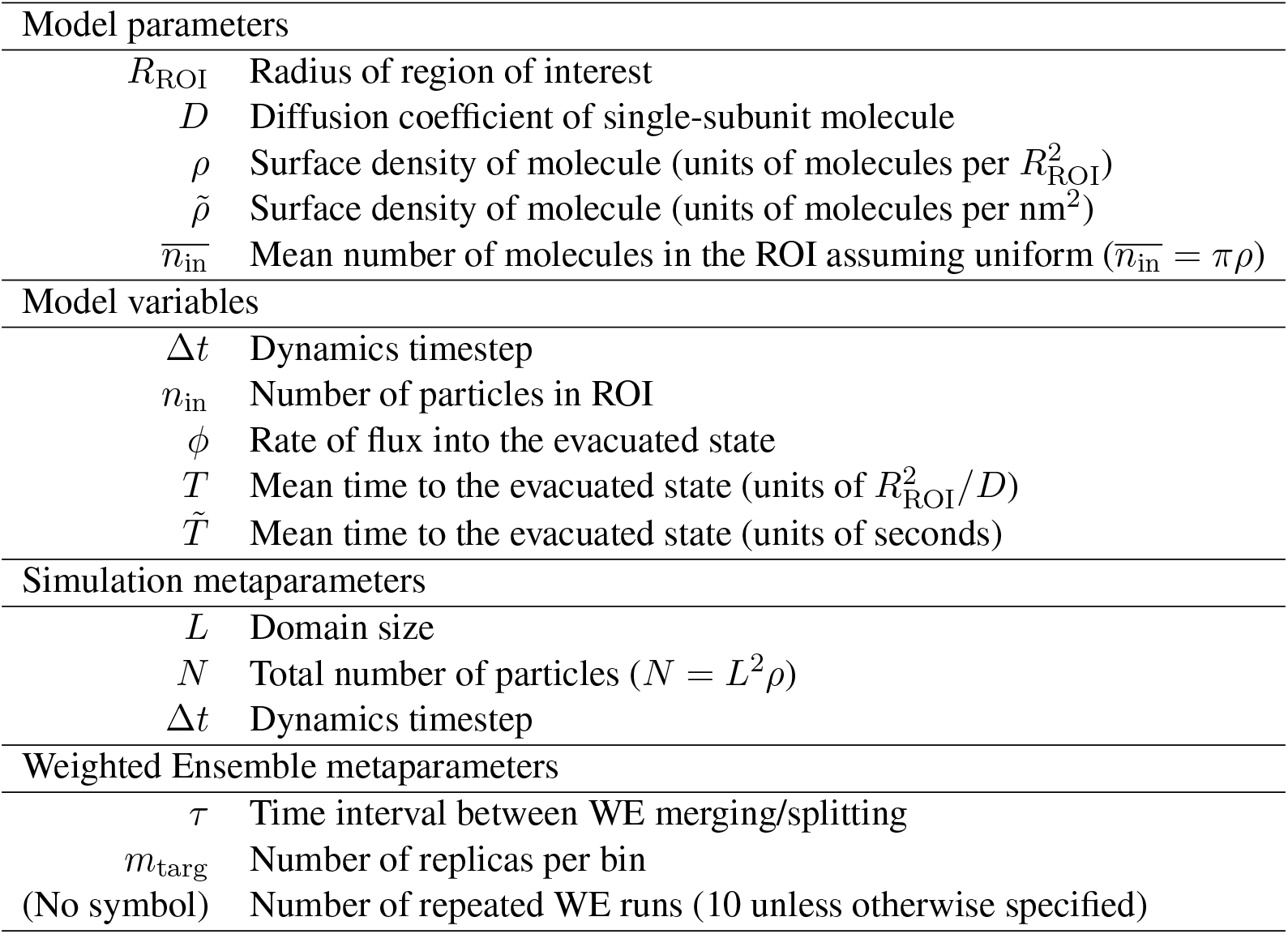
Table of model variables, model parameters, Weighted Ensemble and simulation metaparameters.

| Model parameters |  |
| --- | --- |
| $R_{\text{ROI}}$ | Radius of region of interest |
| $D$ | Diffusion coefficient of single-subunit molecule |
| $\rho$ | Surface density of molecule (units of molecules per $R_{\text{ROI}}^2$) |
| $\tilde{\rho}$ | Surface density of molecule (units of molecules per $\text{nm}^2$) |
| $\bar{n}_{\text{in}}$ | Mean number of molecules in the ROI assuming uniform ($\bar{n}_{\text{in}} = \pi \rho$) |
| Model variables |  |
| $\Delta t$ | Dynamics timestep |
| $n_{\text{in}}$ | Number of particles in ROI |
| $\phi$ | Rate of flux into the evacuated state |
| $T$ | Mean time to the evacuated state (units of $R_{\text{ROI}}^2/D$) |
| $\tilde{T}$ | Mean time to the evacuated state (units of seconds) |
| Simulation metaparameters |  |
| $L$ | Domain size |
| $N$ | Total number of particles ($N = L^2 \rho$) |
| $\Delta t$ | Dynamics timestep |
| Weighted Ensemble metaparameters |  |
| $\tau$ | Time interval between WE merging/splitting |
| $m_{\text{targ}}$ | Number of replicas per bin |
| (No symbol) | Number of repeated WE runs (10 unless otherwise specified) |

### Analytic derivation of *p*_entry_ = 0 FPT distribution

Here we compute the mean first evacuation time from a disc of a collection of molecules undergoing diffusion, when reentry is prohibited (*p*_entry_ = 0). To solve this problem, we first find expressions involving evacuation time for a single molecule placed at a specific location inside the region to be evacuated. We then use that to find the mean first evacuation time for a molecule placed arbitrarily in the domain by averaging its expression across the domain. Lastly, we adapt that expression to systems of *N* non-interacting molecules placed randomly across the domain as an expression of the mean final evacuation time.

#### Single Molecule Evacuation

Let *R* = *R*_ROI_ and *p* = *p*(**r**, *t*|**r**_0_) be the probability density of a single molecule given the molecule’s initial position **r**_0_ = (*r*_0_, *θ*_0_). This probability obeys the two-dimensional diffusion equations

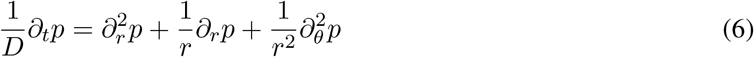

with boundary conditions

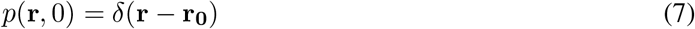

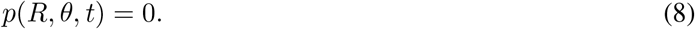

This equation is separable, *p*(*r, θ, t*) = *T*(*t*)Θ(*θ*)*R*(*r*). The separated equations are

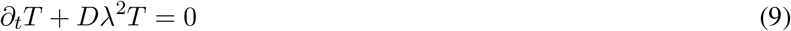

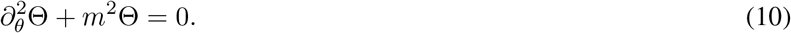

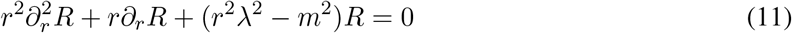

The temporal and angular solutions are

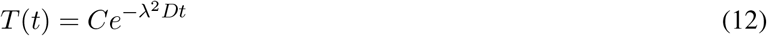

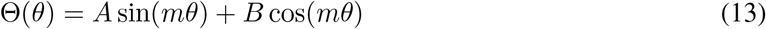

for unknown constants *A,B,C*. The periodicity requires that Θ(*θ*) = Θ(*θ* + 2*π*), which demands *m* be an integer. The radial equation is an example of Bessel’s equation, and given the boundedness of the solution at the origin it is appropriate to limit the possible solutions to Bessel functions of the first kind. This also defines the constant λ to be one of the zeroes of the Bessel function for some corresponding *m*, with *J_m_*(λ_*m,n*_*R*) = 0, where the *m* subscript specifies the order of Bessel function and *n* subscript refers to one of the (infinitely many) zeroes of the function. Combining arbitrary constants our solution may be written generally as

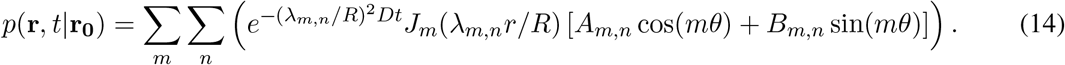

#### Survival Probability and Passage Time Distributions

We introduce the survival probability *G*(*t*), the probability that the molecule remains in the evacuation region at time *t*. This can be expressed in terms of an integral over the position

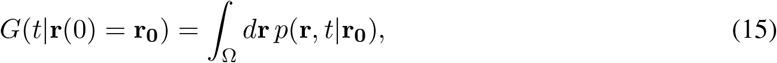

where Ω is the spatial domain of the evacuation of radius *R*. This leads to

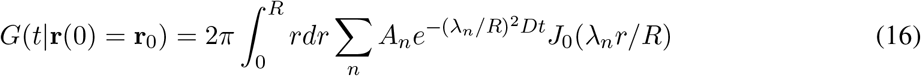

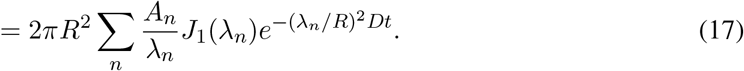

Eq. 16 by recognizing that the integral vanishes when *m* > 0 and relabeling the indices to only count the zeros of *J*_0_. Eq. 17 the property of Bessel functions 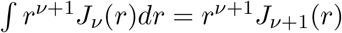 [74].

Using the orthogonality relation

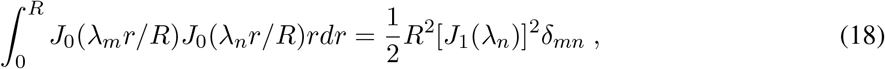

allows us to find the coefficients

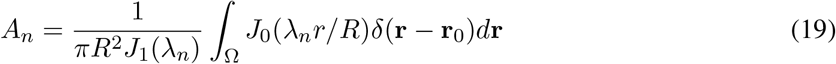

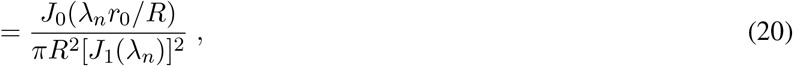

and our final expression for the survival probability *G* is

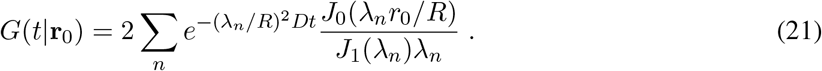

The survival probability can also be written in terms of the distribution of first passage times, *ϱ*(*τ*|**r**_0_), as

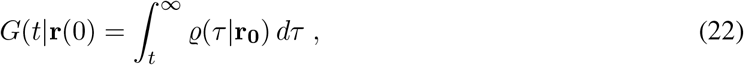

which implies

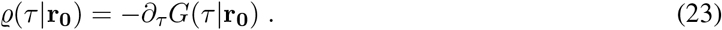

Integration by parts then gives the mean first passage time of a single diffusing molecule in terms of its positional probability distribution,

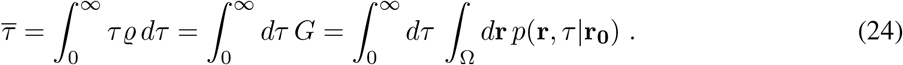

To find the mean first passage time for a molecule randomly placed throughout the domain, rather than at a specific initial condition **r**_0_, we average across all possible initial conditions

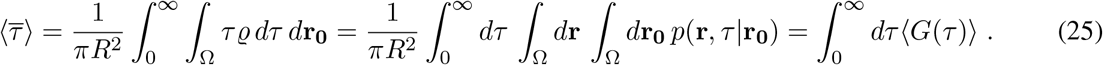

where

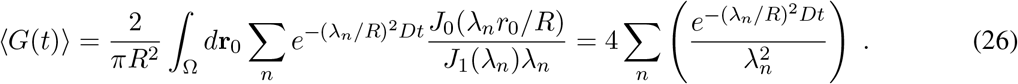

We confirm this result by comparison with brute-force simulation in Fig. S4A.

As an aside, noting that the molecule should always remain in the region at *t* = 0, 〈*G*(*t*)〉 should evaluate to 1 at *t* = 0, implying the following property of the zeroes of the order 0 Bessel function of the first kind:

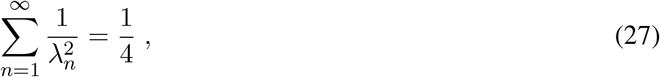

which is an interesting relation that this work confirms numerically.

#### Generalization to *N* Molecules

To generalize this solution for *N* distinct, non-interacting molecules consider the joint probability that all molecules evacuate prior to some time *t*’:

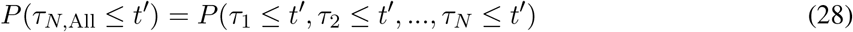

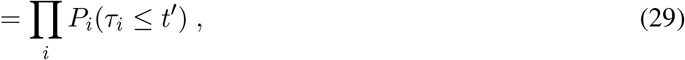

where *P*(*τ_N,AII_* ≤ *t*’) is the probability that the evacuation time for every molecule is less than *t*’. As the molecules are independent and non-interacting, we express it as a product of their individual probabilities of evacuating prior to *t*’, *P_i_*. These *P_i_* are the complement to each molecule’s associated *G*(*t*|**r**_*i*_) given in the previous section,

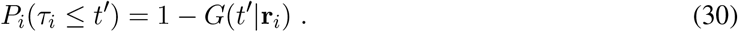

We are interested in the survival probability of the entire system of *N* particles, which we can substitute into Eq. 24 and Eq. 25 to get the mean first passage time of the entire system. We designate the total survival probability *G_N,Any_* (*t*), i.e. the probability that at least one molecule remains inside the evacuation region at some time t, as the complement to the probability of every molecule evacuating.

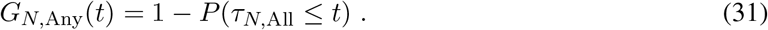

Combining Eq. 31 into Eq. 28 yields

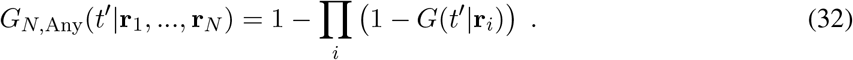

As the molecules are non-interacting, we can use Cov(*G*(*t*’|**r**_1_), *G*(*t*’ |**r**_2_)) = 0 to obtain (*G*(*t*’|**r**_1_)*G*(*t*’|**r**_2_)) = (*G*(*t*’|**r**_1_))(*G*(*t*’ |**r**_2_)) = (*G*(*t*))^2^ and find an expression for the spatial average of the probability at least one molecule remains in the region of interest. The spatial average of *G_N,Any_* can then be written as

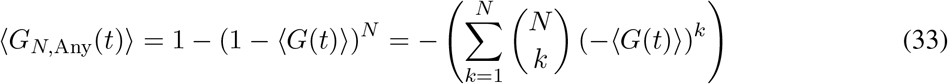

The mean first passage time for the entire system is then given by

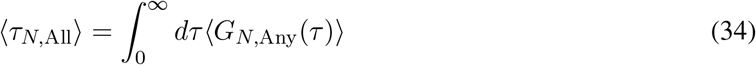

We confirm this result by comparison with brute-force simulation in Fig. S4B.

##### Placement Outside of Evacuation Region

To account for placement outside of the evacuation region, if we have *n* molecules and 〈*τ_m,All_*〉 is the evacuation time for exactly *m* molecules inside the ROI, with *m ≤ N*, we can use the fact that for each molecule, the probability of being placed inside the evacuation region is *p*(**r**_*j*_ ∈ Ω_ROI_) = *A*_ROI_/*A*_Domam_, the ratio of areas of the ROI and the whole domain. The overall evacuation then can be measured as a mean of the possible evacuation times across the Bernoulli trials of successful molecule placement inside the evacuation region,

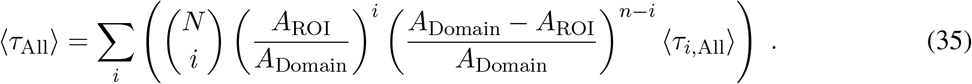

**Figure S4:**
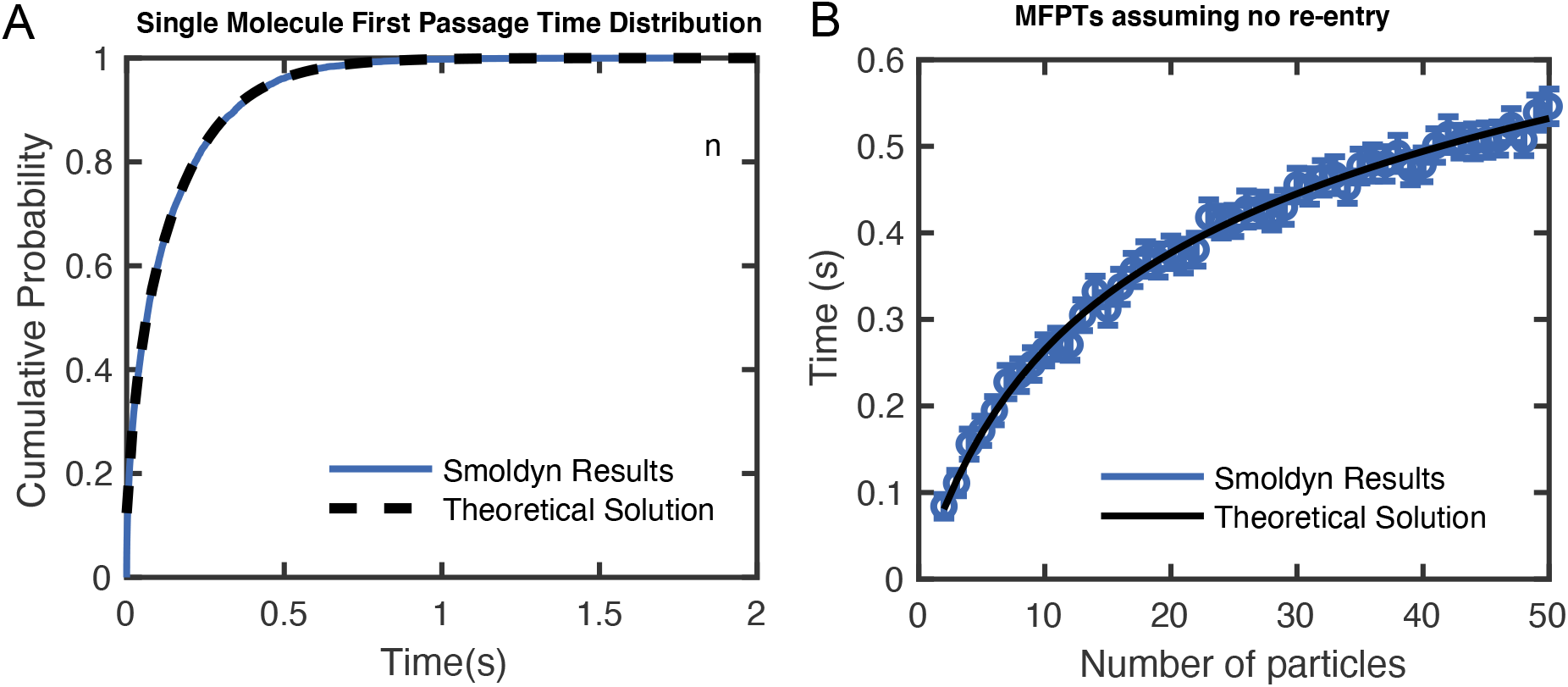
Evaluation of analytical solution for MFPT with *p*_entry_ = 0 shows strong agreement with brute force simulation. (A) Cumulative distribution for the first passage time of a single molecule to leave the ROI from analytical expression in Eq. 26 (black) and Smoldyn results come as the result of 20000 individual simulations (blue). (B) MFPT for multiple molecules inside of the domain, analytically from Eq. 34 (black) and computationally from Smoldyn (blue). In addition to the *N* values shown, an analytic value for N=256 is given in Fig. 6C. Error bars for Smoldyn results are given as the standard error of the mean for 500 Smoldyn simulations per data point.

